# Discovery of Novel Probiotic Species to Improve Infant Health

**DOI:** 10.1101/2025.10.27.684727

**Authors:** Ran Lu, Nakyung Lee, Austin K. Drake, Ryan P. Fulton, Nader Shawky Abutaleb, Mohamed Seleem, David Noah Oakland, Monica P. Garin-Laflam, Mihai Pop, Xin M. Luo

**Affiliations:** Department of Biomedical Sciences and Pathobiology, Virginia-Maryland College of Veterinary Medicine, Virginia Tech, Blacksburg, VA, USA; Department of Computer Science, University of Maryland, College Park, MD, USA; Pediatric Inflammatory Bowel Disease, Carilion Clinic Children’s Hospital, Roanoke, VA, USA; Center for One Health Research, Virginia-Maryland College of Veterinary Medicine, Virginia Tech, Blacksburg, VA, USA

**Keywords:** Breastmilk, formula, infant microbiome, pathogen-specific IgA, probiotics

## Abstract

Selecting an appropriate infant formula can be challenging for parents given the wide range of available options, which may lead to suboptimal choices or frequent switching. Our recent studies in mice demonstrated that the maternal microbiota—particularly *Limosilactobacillus reuteri* present in breast milk—plays a critical role in shaping neonatal immunity by promoting the production of immunoglobulin A (IgA), a key antibody for defense against enteric pathogens. To extend these findings to humans, we analyzed fecal samples from healthy breastfed infants using shotgun metagenomic sequencing combined with whole-bacterial-cell ELISA to assess associations between specific taxa and pathogen-reactive IgA. We found that members of the *Oscillibacter* species (including *Oscillibacter* sp. KLE1728, KLE1745, PC13, PEA192, and *O. valericigenes*) and *Morganella* (e.g., *M. morganii, Morganella* sp. EGD-HP17, and *Morganella* sp. HMSC11D09) were strongly correlated with IgA responses against *Salmonella* and *Shigella*. In contrast, IgA reactive to enterohemorrhagic *Escherichia coli* (EHEC) was associated with other genera, including *Staphylococcus*. Notably, *Lactobacillus* species (*L. bombi, L. kefiri, L. equi*, and *L. rhamnosus*) were consistently linked to elevated IgA responses against *Salmonella, Shigella*, and EHEC. Moreover, levels of anti-*Salmonella* and anti-*Shigella* IgA were positively correlated with the use of infant formula containing prebiotic fibers or human milk oligosaccharides (5-HMO). Collectively, these findings highlight novel commensal taxa with potential as probiotic candidates to guide the development of next-generation infant formulas aimed at enhancing mucosal immunity and supporting infant health.

**Importance:** Early-life nutrition shapes immune maturation, yet the microbial factors driving mucosal antibody responses remain poorly defined. Through integrated metagenomic and IgA profiling, this study identifies commensal taxa, particularly *Lactobacillus, Oscillibacter*, and *Morganella*, associated with pathogen-specific IgA in infants. These taxa may enhance mucosal defense against enteric pathogens, revealing key microbial contributors to early immune development. The findings provide a foundation for designing next-generation infant formulas that leverage targeted probiotics or prebiotics to promote protective IgA responses and gut health.

## Introduction

Breastmilk is believed to provide not just nutrition but also immune, metabolic, and neurodevelopmental health. Human milk delivers a complex mixture of immunoglobulins, cytokines, and maternal immune cells that confer immediate protection against pathogens while promoting maturation of the neonatal immune system ^1^. Among these, secretory IgA (SIgA) and other bioactive components play central roles in shaping mucosal immunity and regulating inflammatory responses in the infant gut ^1; 2^. Breastfeeding also promotes the establishment of a gut microbiota enriched in *Bifidobacterium* and other commensals that are critical for immune education and intestinal barrier development ^3^. Epidemiological studies consistently show that breastfed infants have lower risks of respiratory and gastrointestinal infections and reduced incidence of chronic diseases such as asthma, obesity, and type 2 diabetes later in life ^2; 4^. Furthermore, breastmilk–derived fatty acids and neurotrophic factors have been linked to enhanced cognitive outcomes ^5; 6^. Collectively, these findings underscore the multifaceted benefits of breastfeeding and highlight the importance of identifying breastmilk–derived microbes and immune modulators that could inform the design of improved infant formulas.

In addition to providing food for the gut microbiota, breastmilk also acts as a source of maternal microbes that seed the infant gut microbiome. Infants deprived of this early inoculation exhibit distinct microbial communities, often characterized by reduced colonization with beneficial taxa such as *Bifidobacterium longum subsp. infantis* and *Lactobacillus* species, and greater abundance of potentially pro-inflammatory bacteria including *Clostridium* and *Enterobacteriaceae* ^3; 7; 8^. These alterations have been associated with impaired fecal IgA production, weakened mucosal defenses, and increased risks of infections, allergic diseases, and chronic inflammation ^9; 10^. In addition, human milk oligosaccharides (HMOs) selectively promote the growth of beneficial bacteria, creating a synergistic relationship between milk glycans and probiotics that further strengthens host–microbe interactions during a critical developmental window ^11^. Formula-fed infants lacking HMOs and breastmilk–derived microbes may therefore experience suboptimal microbial–immune crosstalk.

IgA is the most abundant antibody isotype in humans and a cornerstone of mucosal defense. In the intestine, SIgA is actively transported across epithelial cells into the lumen, where it orchestrates host–microbe interactions ^12^. Microbial colonization after birth provides key signals for gut-associated lymphoid tissue (GALT) maturation and the development of IgA-secreting plasma cells ^13^. Commensals stimulate IgA production through both T cell–dependent pathways—characterized by germinal center formation, class switching, and affinity maturation—and T cell–independent pathways, which generate low-affinity, polyreactive IgA ^14; 15^. Once secreted, IgA mediates immune exclusion, promotes microbial agglutination, modulates bacterial gene expression, and contributes to symbiosis. Importantly, SIgA can selectively coat commensals, shaping microbial spatial organization and abundance without inducing inflammation ^16^. Our recent studies in mice have shown that the maternal microbiota, particularly *Limosilactobacillus reuteri* found in breastmilk, plays a critical role in shaping the neonatal immune system by promoting the production of IgA ^17^.

Disruption of this IgA–microbiota axis can have significant consequences. Individuals with selective IgA deficiency exhibit altered microbiota with increased prevalence of pathobionts, reduced diversity, and heightened susceptibility to infections, allergies, and autoimmunity ^18; 19^. Dysregulated IgA responses have also been implicated in inflammatory bowel disease, celiac disease, and metabolic disorders, where aberrant IgA coating of commensals correlates with disease activity ^20^. These findings suggest that IgA is not only a biomarker of mucosal immune status but also an active mediator of microbial homeostasis.

Therefore, breastfeeding delivers protective microbes and IgA-inducing factors essential for mucosal immune development. Infants who are not breastfed may miss these cues, predisposing them to dysbiosis and immune dysregulation. Current formulas, although nutritionally adequate, lack the immunomodulatory functions of breastmilk, particularly IgA-inducing commensals. Identifying and incorporating these components into formula represents a critical step toward replicating the immune-training effects of breastfeeding and reducing long-term risks of infection and immune-mediated disease in non-breastfed infants. Specifically, the identification of neonatal IgA-inducing commensal bacterial strains will guide formula selection as well as future formula design to include such probiotic strains or include the prebiotics capable of promoting the growth and/or gut colonization of these probiotic strains. In addition, by assessing the fecal microbiome of breastfed infants and establishing its relationship with mucosal IgA response, we can discover additional commensal bacterial strains promoted by breastmilk that are associated with better mucosal immune defense. These newly identified gut bacteria may become novel probiotics to be added to the infant formula.

## Materials and Methods

### Study Design and Participant Enrollment

This prospective longitudinal study enrolled 9 healthy, breastfed or breastmilk-formula mixed fed infants from three pediatric clinics, including general pediatric offices and one gastrointestinal specialty clinic. The study protocol was approved by the Institutional Review Board (IRB) at Carilion Clinic with IRB #21-1327. Written informed consent was obtained from the parents of all participants prior to enrollment. Inclusion criteria for participants included exclusive breastfeeding, breastmilk and formula mixed feeding, no history of bacterial gastrointestinal infection, no receipt of an oral bacterial vaccination, and no prior antibiotic use. A physical examination was performed to determine eligibility.

### Sample Collection and Handling

Fecal samples were collected longitudinally from each enrolled infant at 1, 2, 4, and 6 months of age. Samples collected during a clinic visit were immediately transferred to a provided container and frozen at −20°C. For samples not provided during a visit, parents were given a stool collection kit and instructed to collect the sample at home, place it in the freezer immediately, and return it to the clinic on their next visit. All samples were transported to Virginia Tech at −20°C and stored at −80°C until further analysis.

### Laboratory Analyses

All laboratory procedures were performed in a Biosafety Level 2 (BSL-2) laboratory at Virginia Tech. Total fecal IgA was determined with a commercial kit (BioVendor). Bacteria-specific IgA was assessed for reactivity to *Salmonella typhimurium, Shigella flexneri* (ATCC 9199) and enterohemorrhagic *E. coli* (EHEC) by a previously developed whole-bacteria ELISA ^17^. Briefly, *Salmonella, Shigella* and EHEC cells were fixed with 1% formalin for 2 hours at room temperature. Approximately 10^8^ colony-forming units of fixed bacterial cells in 100 μL of coating buffer (50 mM sodium carbonate) were added into each well of a Nunc Maxisorp plate (Thermo Fisher, Catalog number 44-2404-21) and incubated overnight at 4°C. The plate was blocked with 4% non-fat milk in PBS-T (phosphate-buffered saline with 0.05% Tween-20) for 2 hours at room temperature. Subsequently, 50 μL of a serially diluted antiserum (positive control) or samples were added to the wells for a 1-hour incubation at room temperature. *E. coli* O157:H7 antiserum (Seracare) was diluted starting from 1:10000. *Shigella* antisera (Fisher Scientific) was diluted staring from1:1000. *Salmonella* H antiserum I (1:80000), O antiserum Factors 4,5 (1:80000), and O antiserum Group D1 (1:64) (BD DifcoTM) were combined prior to serial dilution. The wells were then incubated for 1 hour at room temperature with a horseradish peroxidase-conjugated anti-human IgA detection antibody (Southern Biotech). Super AquaBlue ELISA substrate (Thermo Fisher) was used for color development. The plate’s optical density was read at OD405 using a SpectraMax M5 (Molecular Devices). Autoantigen microarrays were performed by GeneCopoeia, Inc. (Rockville, MD).

### Microbiome and Bioinformatic Analyses

To prepare bacterial DNA samples for microbiome analysis, 100 mg of frozen fecal material was placed in Fast Prep Lysing Matrix D tubes containing ceramic beads (MP Biomedicals) and incubated in 1 mL of PBS on ice for 1 hour. Fecal pellets were homogenized by bead beating for 5 seconds and then centrifuged (50 × *g*, 15 minutes, 4°C) to remove large particles. Fecal bacteria in the supernatants were collected, washed with 1 mL of PBS containing 1% (w/v) bovine serum albumin (BSA), and centrifuged for 5 minutes at 8,000 × *g*, 4°C. Total bacterial DNA was isolated using a commercial kit (Qiagen) then submitted to the Fralin Life Sciences Institute Genomics Sequencing Center at Virginia Tech for library preparation and metagenomic sequencing. DNA-seq libraries were constructed using Kapa Hyper Prep Library Kit from Roche (Catalog No. 7962347001). Library preparation: input DNA was quantitated using a qubit 3.0.

Samples were then sheared using a Covaris M220. End Repair and A tailing were performed on the roughly 750 ng input DNA. Adapter ligation and barcoding were performed followed by a bead clean up and PCR (3 cycles). Agilent Tapestation was used to visualize the final libraries which were quantitated using Quant-iT dsDNA HS Kit (Invitrogen). Libraries were then normalized and pooled and sequenced on a NovaSeq 6000 S1 300 cycle kit (150bp Paired End Reads). The Illumina NovaSeq Control Software v1.7 was used to provide the management and execution of the NovaSeq 6000 and to generate BCL files. The BCL files were converted to FASTQ files, adapters trimmed and demultiplexed using bcl2fastq Conversion Software v2.20. At the University of Maryland, sequencing data was verified with fastqc and trimmed with bbduk ^21^. After quality assessment, reads were mapped against hg38 reference genome with bowtie to filter out reads associated with human. Non-human reads (i.e. unmapped reads) were extracted using SAMtools and then assembled using MEGAHIT (max k-mer size 141). These assemblies were then classified by taxonomy using Kraken2 to identify the presence and abundance of microbial organisms. Finally, the results were used to measure the relationship between taxonomy classification and IgA information through pearson correlation. Sequencing data are available from the NCBI database with the accession number PRJNA1338259.

## Results and Discussion

Understanding bacterial strain-specific differences is crucial for the selection of appropriate pre/probiotics for formula enrichment. With many products on the market, it is challenging to design a robust clinical study to identify the best pre/probiotics that would enhance the neonatal mucosal antibody response against infections. We investigate the correlation between infant microbiome and mucosal immunity. The rationale is that the infant microbiome reflects the input provided by either breastmilk or any baby formula with an appropriate formulation of pre/probiotics. Practically, we just need infant stools from which both the gut microbiome and mucosal IgA can be quantified. Correlation analyses can then be performed to reveal specific gut bacteria that are associated with an enhanced mucosal immune response, thus guiding probiotic formulation to enhance infant health.

We longitudinally profiled pathogen-specific fecal IgA in nine infants across the first six months of life. While most infants displayed little or no IgA reactivity to *Salmonella, Shigella*, or EHEC, a subset exhibited striking responses. For example, Infant 107 produced broadly polyreactive IgA against all three pathogens, whereas infant 102 mounted selective responses, with strong anti-*Salmonella* and anti-*Shigella* but weak anti-EHEC IgA. Infant 110 and 109 showed unusually high anti-*Salmonella* IgA as early as one month (Figure 1). These data highlight marked inter-individual heterogeneity in fecal IgA development, even in predominantly breastfed infants who receive maternal microbes and secretory IgA (SIgA). A likely explanation is that breastmilk-associated microbes stimulate early IgA responses in infants, but these tend to be of relatively low affinity, consistent with the polyreactive nature observed in some infants. In contrast, breastmilk and formula mixed fed infants (infant 102,107,109 and 110) showed higher IgA responses against *Salmonella* and *Shigella* compared to infants that are breastfed exclusively (Figure 2). This is potentially driven by human milk oligosaccharide (HMO) mimetics and prebiotics included in their formulas that potentially promote more targeted B cell activation to produce *de novo* IgA. Such variation likely reflects a combination of factors including differences in maternal microbes, infants early microbial colonization dynamics ^8; 22^, maternal antibody repertoires ^23^, infant genetic background ^24^ and timing of immune maturation ^10^. Importantly, this data suggests that the source and type of microbial exposure—whether through breastmilk microbiota or prebiotic-enriched formula—can shape the affinity and breadth of neonatal IgA responses, with implications for protection against enteric pathogens during early life.

**Figure 1.**
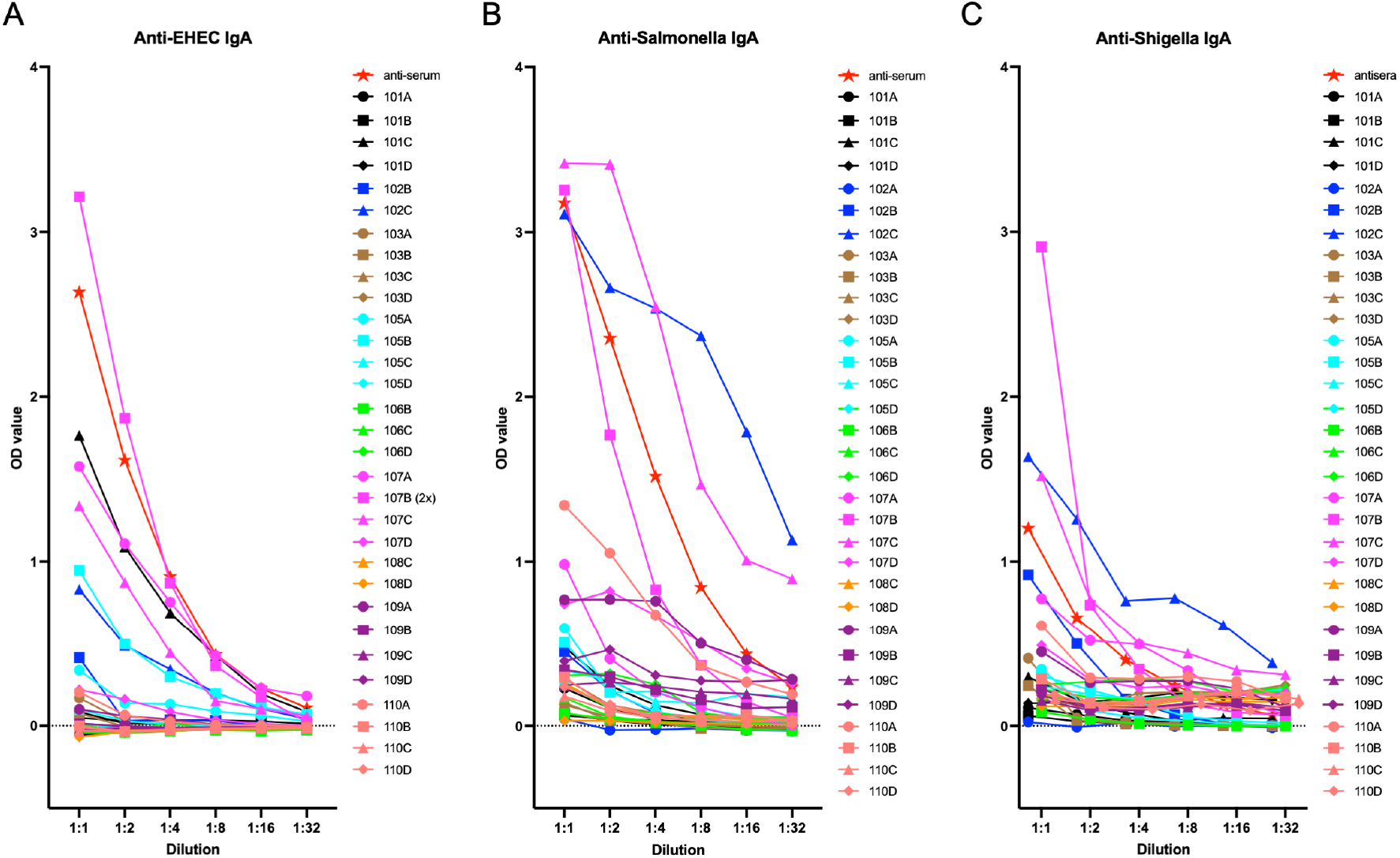
Detection of anti-pathogen fecal IgA in infants by bacterial whole-cell ELISA. Fecal samples from infants were analyzed for IgA reactivity against (A) enterohemorrhagic *E. coli* (EHEC), (B) *Salmonella*, and (C) *Shigella* using bacterial whole-cell ELISA. Serial dilutions of fecal samples (1:1 to 1:32) were applied to pathogen-coated plates, and optical density (OD) values were recorded to measure pathogen-reactive IgA binding. Each curve represents an individual sample. Commercially available anti-sera were used as the positive controls.

**Figure 2.**
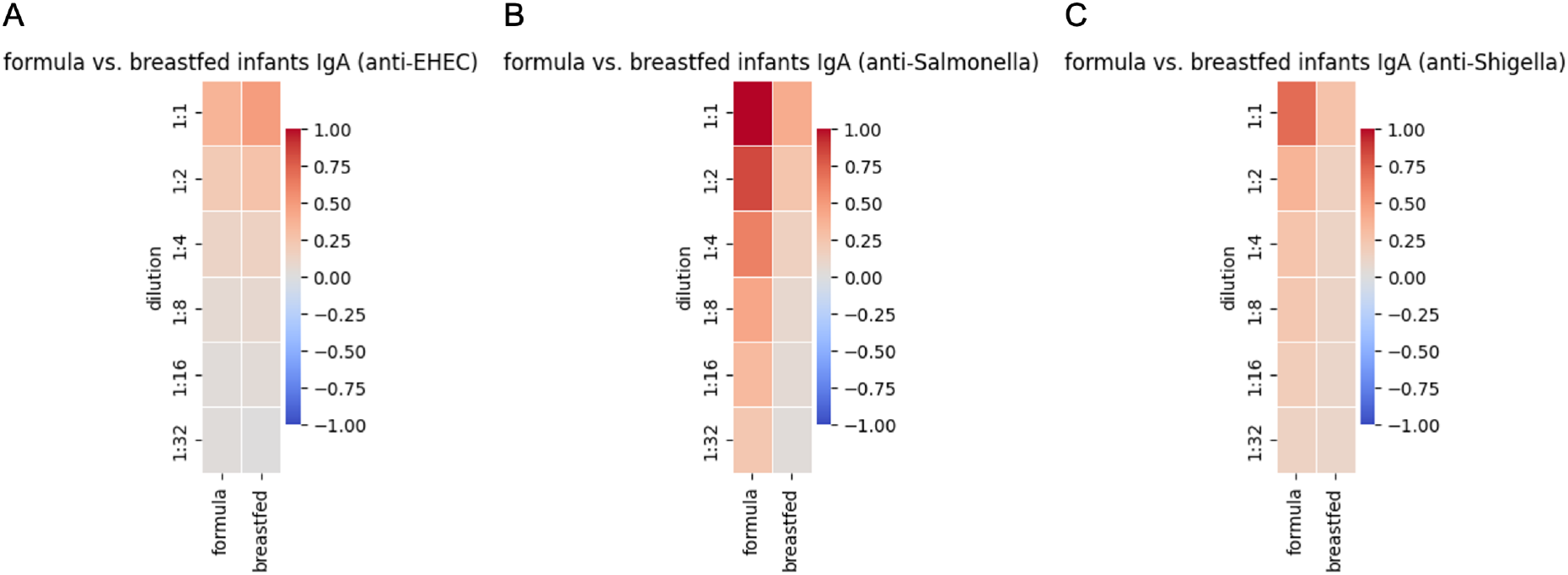
Comparison of pathogen-reactive IgA responses between formula-fed and breastfed infants. Fecal IgA reactivities against (A) EHEC, (B) *Salmonella*, and (C) *Shigella* were assessed by bacterial whole-cell ELISA and compared between formula-fed (n=4) and exclusively breastfed (n=5) infants.

Formula exposure was associated with increased fecal anti-*Salmonella* and anti-*Shigella* IgA, whereas no effect was observed for anti-EHEC IgA. While preliminary, this finding is intriguing and may reflect the combined influence of formula components (e.g., HMO or added prebiotics) ^25; 26^ and microbial alterations induced by mixed feeding ^3^. HMOs and prebiotics are known to enrich for specific taxa capable of modulating IgA responses ^27^, and formula feeding is often linked to increased microbial diversity ^7^, which may enhance antigen exposure and mucosal immune activation. The absence of an effect on EHEC-specific IgA further suggests that pathogen-specific IgA responses are regulated by distinct microbial and dietary cues.

Microbiome analyses revealed distinct microbial signatures associated with pathogen-specific IgA responses. Abundance of commensals such as *Oscillibacter, Fusobacterium, Coprobacillus* and *Morganella* correlated strongly with all three anti-*Salmonella*, anti-*Shigella and anti-EHEC* IgAs (Figure 3), suggesting that these commensals may provide shared antigenic or metabolic cues that elicit cross-reactive IgA against these pathogens ^13; 28; 29^. Commensals such as *Klebsiella, Clostridium, Hungatella, Atlantibacter, Megasphaera* and *Lactobacillus* exhibiting similar correlation patten with both anti-Salmonella and anti-Shigella IgA reactivities.

**Figure 3.**
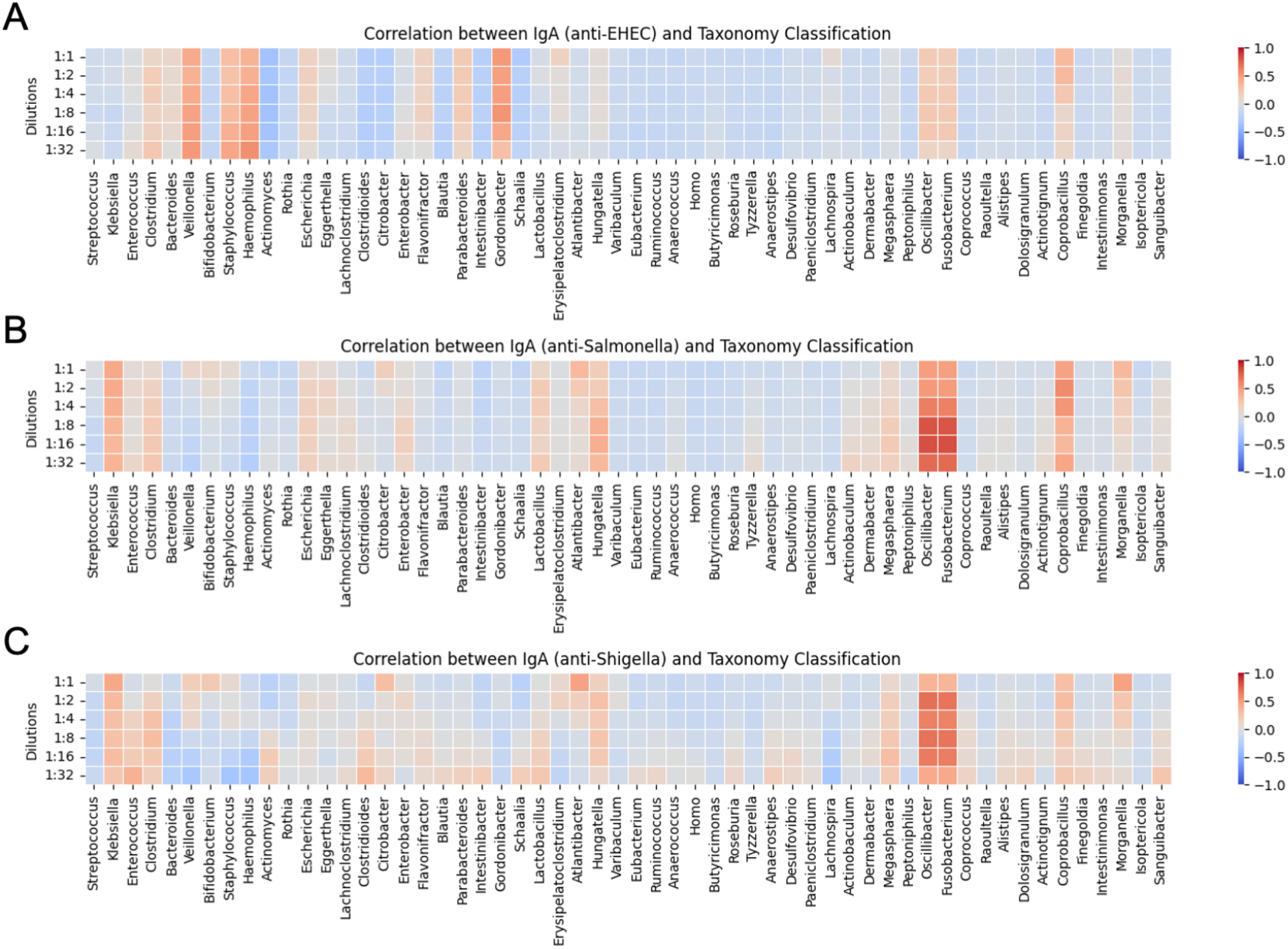
Correlations between pathogen-reactive IgA responses and gut microbiome composition at the genus level. Bacterial DNA was extracted from infant fecal samples and analyzed by shotgun metagenomic sequencing to profile gut microbiome at the genus level. Correlations are shown between the relative abundance of bacterial genera and fecal IgA against (A) EHEC, (B) *Salmonella*, and (C) *Shigella*.

Interestingly, anti-EHEC IgA showed a distinct correlation with commensals involving *Staphylococcus, Veillonella, Haemophilus and Gordonibacter*, underscoring pathogen-specificity in microbiota–immune interactions. While *Fusobacterium* species have been implicated in immunomodulatory interactions with the host, they are also recognized as opportunistic pathogens ^30; 31^. Their dual nature underscores the need for careful risk–benefit evaluation before considering them for probiotic development, and accordingly we did not include them among our candidate species. *Klebsiella* is likewise frequently associated with pathogenicity and, in many contexts, is considered a more significant health concern than *Fusobacterium*. At the same time, *Klebsiella* species have also been detected in breastmilk and the infant gut microbiome, suggesting that not all members of this genus are uniformly pathogenic ^8; 32^. Nonetheless, given its strong clinical association with disease, *Klebsiella* was excluded from consideration in our study.

At the species level, *Coprobacillus sp*. 8-2-548FAA and multiple *Lactobacillus* specie*s (L. Bombi, L. rahmosus, L. kefiri and L. equi*) correlate to all three pathogen specific IgA. Multiple *Oscillibacter* (e.g., *Oscillibacter* sp. KLE1728, KLE1745, PC13, PEA192, and *O. valericigenes*), *Morganella* (e.g., *Morganella morganii, Morganella sp*. EGD-HP17 and *Morganella sp*. HMSC11D09) were linked to anti-*Salmonella* and anti-*Shigella* IgA, while anti-EHEC IgA correlates with more complex taxa composition in *Veillonella* genus (Figure 4).

**Figure 4.**
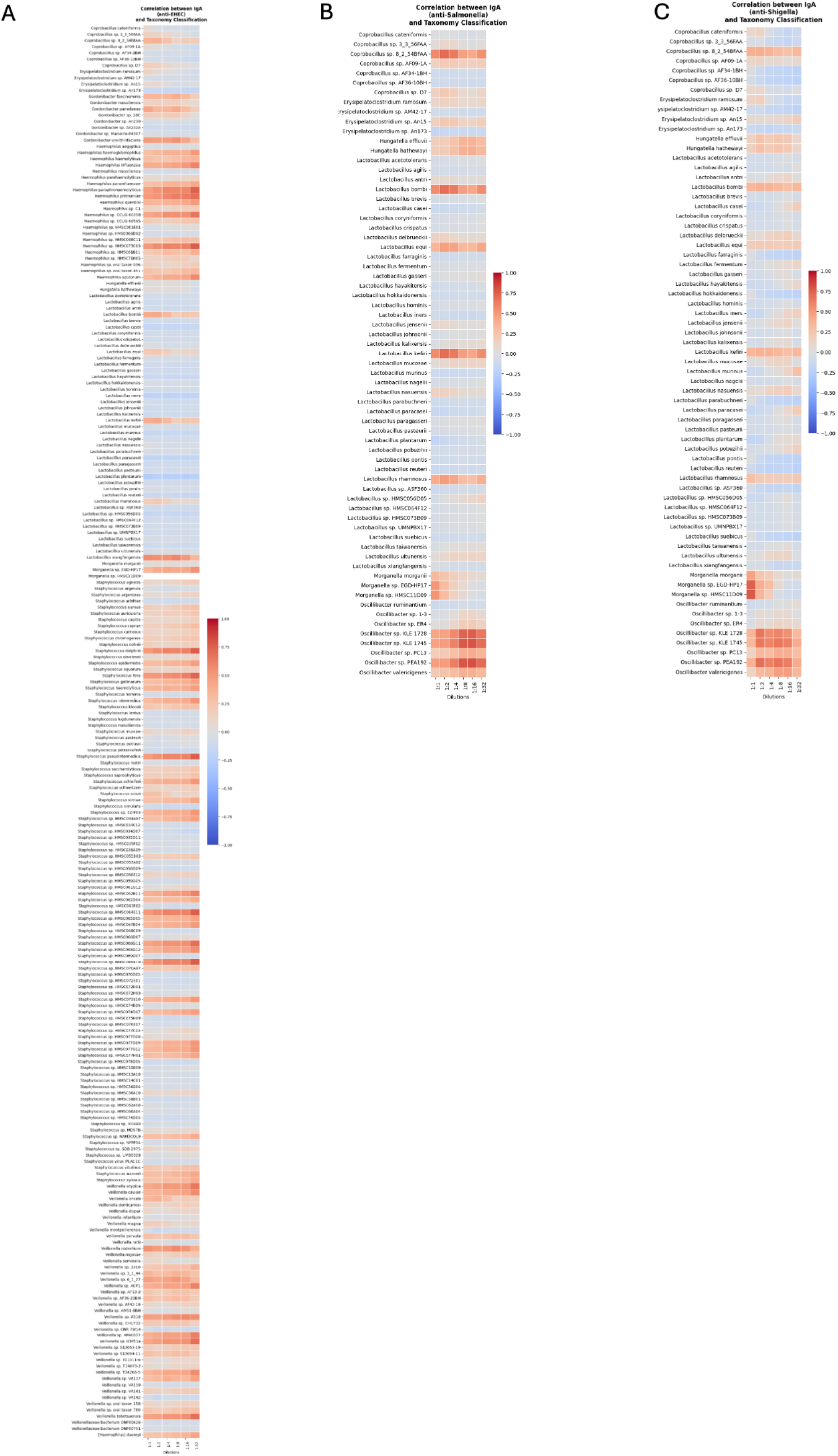
Correlations between pathogen-reactive fecal IgA responses and gut microbiome composition at the species level. Fecal samples from infants were subjected to shotgun metagenomic sequencing to characterize gut microbial composition at the species level. Correlative analyses were performed between the relative abundances of bacterial species and fecal IgA reactivity against (A) EHEC, (B) *Salmonella*, and (C) *Shigella*.

For infants who cannot be exclusively breastfed, reduced exposure to breast milk microbes represent a major immunological gap. Our findings suggest *Lactobacillus* spp. *(L. Bombi, L. rahmosus, L. kefiri and L. equi*), *Oscillibacter* (e.g., *Oscillibacter* sp. KLE1728, KLE1745, PC13, PEA192, and *O. valericigenes*) and *Morganella* (e.g., *Morganella morganii, Morganella sp*. EGD-HP17 and *Morganella sp*. HMSC11D09) as candidate IgA-inducing commensals that could be harnessed in probiotic-fortified formulas. *Oscillibacter* species are emerging as producers of valeric acid and other metabolites that enhance IgA and barrier integrity ^33; 34^, while *Lactobacillus* strains are well-established IgA inducers that also promote immune tolerance and restrict pathogen colonization ^17; 35-41^. Interestingly, *Morganella*—traditionally studied as an opportunistic pathogen—has recently gained attention for its potential immunomodulatory roles within the gut microbiome ^30^. Several studies suggest that *Morganella morganii* and related taxa can engage host pattern recognition receptors (e.g., TLR4, TLR5) through their lipopolysaccharides and flagellin, thereby stimulating innate immune responses ^42^. Moreover, metagenomic analyses have found correlations between *Morganella* abundance and heightened IgA coating in fecal IgA-seq studies, suggesting that these bacteria are frequent targets of adaptive immune recognition ^43^. While their dual identity as both commensal residents and opportunistic pathogens warrants careful safety evaluation, selected *Morganella* strains may harbor immunostimulatory properties that could be harnessed for controlled IgA induction and mucosal immune education in early life.

Incorporating such microbes into infant formulas, ideally in combination with HMOs or targeted prebiotics, offers a rational strategy to mimic the dual functions of breastmilk: passive protection via antibodies and active immune education via microbes. A range of probiotic strains has been incorporated into commercially available infant formulas, most notably *Bifidobacterium animalis* subsp. *lactis* (e.g., BB-12, HN019), *Bifidobacterium breve* (M-16V), *Bifidobacterium infantis, Lactobacillus rhamnosus* GG, and *Lactobacillus reuteri* DSM 17938, either alone or in synbiotic formulations with prebiotic oligosaccharides. Clinical investigations consistently demonstrate that such products are safe and support growth outcomes equivalent to those of standard formulas ^44; 45^. Reported benefits include reduced incidence of infectious diarrhea and atopic dermatitis, improved stool characteristics, and partial restoration of a gut microbiota profile resembling that of breastfed infants ^46; 47^. However, most clinical endpoints remain descriptive, and mechanistic evidence delineating how specific strains influence infant immune development is still limited. Simply adding perceived “beneficial bacteria” is unlikely to replicate the complex immune-modulating properties of breastmilk. Instead, strain selection should be guided by rigorous functional data, including IgA-inducing capacity, safety, and ecological compatibility with the infant gut. The present study extends this literature by providing IgA-correlating commensals that may inform the rational design of next-generation probiotic formulas. Such targeted probiotic fortification could help recalibrate mucosal immunity in formula-fed infants, reducing risks of infection, allergy, and immune dysregulation.

Our findings provide a framework for the rational design of infant formula supplemented with targeted probiotic candidates. While fecal IgA in breastfed infants reflects both maternal SIgA and infant *de novo* synthesis modulated by breastmilk microbes ^48^, the latter becomes increasingly dominant during the first year of life ^49; 50^. Our results highlight that specific commensal, particularly *Lactobacillus (L. Bombi, L. rahmosus, L. kefiri and L. equi*) and *Oscillibacter* (*Oscillibacter* sp. KLE1728, KLE1745, PC13, PEA192, and *O. valericigenes*) species, are consistently associated with pathogen-specific IgA responses. This suggests that formula enriched with such probiotic species could recapitulate aspects of the protective functions of breastmilk by fostering finely tuned IgA responses. One limitation of our work is that the associations we describe are correlational and cannot establish causality. While the data highlight potential links between commensal bacteria and pathogen-specific IgAs, establishing a causal relationship would require well-controlled clinical trials testing defined probiotic formulations. Nevertheless, this does not undermine the significance of our findings, which provide important insights into host–microbe interactions and generate testable hypotheses that can guide future interventional studies.

Together, these findings point toward the possibility of leveraging naturally occurring commensals as probiotic additives in infant formula to promote more effective mucosal immunity during early life. Future work will need to pinpoint the specific strains of commensals that most effectively shape pathogen-targeted IgA responses. Achieving this level of precision could open the door to next-generation infant formulas designed for personalized immune support during early life.

## Acknowledgment

This work was funded by an iTHRIV subaward from University of Virginia with NIH award number UL1TR003015.

## References

1. Walker A. Breast milk as the gold standard for protective nutrients. J Pediatr. 2010;156(2 Suppl):S3–7. 10.1016/j.jpeds.2009.11.021

2. Duijts L, Jaddoe VW, Hofman A, Moll HA. Prolonged and exclusive breastfeeding reduces the risk of infectious diseases in infancy. Pediatrics. 2010;126(1):e18– 25. 10.1542/peds.2008-3256

3. Pannaraj PS et al. Association between breast milk bacterial communities and establishment and development of the infant gut microbiome. JAMA Pediatr. 2017;171(7):647–654. 10.1001/jamapediatrics.2017.0378

4. Victora CG et al. Breastfeeding in the 21st century: Epidemiology, mechanisms, and lifelong effect. Lancet. 2016;387(10017):475–490. 10.1016/S0140-6736(15)01024-7

5. Horta BL, Loret de Mola C, Victora CG. Breastfeeding and intelligence: A systematic review and meta-analysis. Acta Paediatr. 2015;104(467):14–19. 10.1111/apa.13139

6. Hou L et al. Impact of the duration of breastfeeding on the intelligence of children: A systematic review with network meta-analysis. Breastfeed Med. 2021;16(9):687–696. 10.1089/bfm.2020.0364

7. Azad MB et al. Gut microbiota of healthy canadian infants: Profiles by mode of delivery and infant diet at 4 months. CMAJ. 2013;185(5):385–394. 10.1503/cmaj.121189

8. Backhed F et al. Dynamics and stabilization of the human gut microbiome during the first year of life. Cell Host Microbe. 2015;17(6):852. 10.1016/j.chom.2015.05.012

9. Henrick BM et al. Bifidobacteria-mediated immune system imprinting early in life. Cell. 2021;184(15):3884–3898 e3811. 10.1016/j.cell.2021.05.030

10. Rogier EW, Frantz AL, Bruno ME, Kaetzel CS. Secretory iga is concentrated in the outer layer of colonic mucus along with gut bacteria. Pathogens. 2014;3(2):390–403. 10.3390/pathogens3020390

11. Marcobal A et al. Consumption of human milk oligosaccharides by gut-related microbes. J Agric Food Chem. 2010;58(9):5334–5340. 10.1021/jf9044205

12. Pabst O, Slack E. Iga and the intestinal microbiota: The importance of being specific. Mucosal Immunol. 2020;13(1):12–21. 10.1038/s41385-019-0227-4

13. Macpherson AJ, Geuking MB, McCoy KD. Immune responses that adapt the intestinal mucosa to commensal intestinal bacteria. Immunology. 2005;115(2):153–162. 10.1111/j.1365-2567.2005.02159.x

14. Fagarasan S, Kawamoto S, Kanagawa O, Suzuki K. Adaptive immune regulation in the gut: T cell-dependent and t cell-independent iga synthesis. Annu Rev Immunol. 2010;28(243–273. 10.1146/annurev-immunol-030409-101314

15. Kubinak JL, Round JL. Do antibodies select a healthy microbiota? Nat Rev Immunol. 2016;16(12):767–774. 10.1038/nri.2016.114

16. Palm NW et al. Immunoglobulin a coating identifies colitogenic bacteria in inflammatory bowel disease. Cell. 2014;158(5):1000–1010. 10.1016/j.cell.2014.08.006

17. Mu Q et al. Regulation of neonatal iga production by the maternal microbiota. Proc Natl Acad Sci U S A. 2021;118(9):10.1073/pnas.2015691118

18. Fadlallah J et al. Microbial ecology perturbation in human iga deficiency. Sci Transl Med. 2018;10(439):10.1126/scitranslmed.aan1217

19. Zhang J, van Oostrom D, Li J, Savelkoul HFJ. Innate mechanisms in selective iga deficiency. Front Immunol. 2021;12(649112. 10.3389/fimmu.2021.649112

20. Viladomiu M et al. Iga-coated e. Coli enriched in crohn’s disease spondyloarthritis promote t(h)17-dependent inflammation. Sci Transl Med. 2017;9(376):10.1126/scitranslmed.aaf9655

21. Bolger AM, Lohse M, Usadel B. Trimmomatic: A flexible trimmer for illumina sequence data. Bioinformatics. 2014;30(15):2114–2120. 10.1093/bioinformatics/btu170

22. Yatsunenko T et al. Human gut microbiome viewed across age and geography. Nature. 2012;486(7402):222–227. 10.1038/nature11053

23. Brandtzaeg P. The mucosal immune system and its integration with the mammary glands. J Pediatr. 2010;156(2 Suppl):S8–15. 10.1016/j.jpeds.2009.11.014

24. Olsson PG, Hammarstrom L, Cox DW, Smith CI. Involvement of both hla and ig heavy chain haplotypes in human iga deficiency. Immunogenetics. 1992;36(6):389–395. 10.1007/BF00218046

25. Chichlowski M, De Lartigue G, German JB, Raybould HE, Mills DA. Bifidobacteria isolated from infants and cultured on human milk oligosaccharides affect intestinal epithelial function. J Pediatr Gastroenterol Nutr. 2012;55(3):321– 327. 10.1097/MPG.0b013e31824fb899

26. Borewicz K et al. Correlating infant fecal microbiota composition and human milk oligosaccharide consumption by microbiota of 1-month-old breastfed infants. Mol Nutr Food Res. 2019;63(13):e1801214. 10.1002/mnfr.201801214

27. Underwood MA, German JB, Lebrilla CB, Mills DA. Bifidobacterium longum subspecies infantis: Champion colonizer of the infant gut. Pediatr Res. 2015;77(1-2):229–235. 10.1038/pr.2014.156

28. Kau AL, Ahern PP, Griffin NW, Goodman AL, Gordon JI. Human nutrition, the gut microbiome and the immune system. Nature. 2011;474(7351):327–336. 10.1038/nature10213

29. Belkaid Y, Hand TW. Role of the microbiota in immunity and inflammation. Cell. 2014;157(1):121–141. 10.1016/j.cell.2014.03.011

30. Shroff KE, Meslin K, Cebra JJ. Commensal enteric bacteria engender a self-limiting humoral mucosal immune response while permanently colonizing the gut. Infect Immun. 1995;63(10):3904–3913. 10.1128/iai.63.10.3904-3913.1995

31. Talham GL, Jiang HQ, Bos NA, Cebra JJ. Segmented filamentous bacteria are potent stimuli of a physiologically normal state of the murine gut mucosal immune system. Infect Immun. 1999;67(4):1992–2000. 10.1128/IAI.67.4.1992-2000.1999

32. Jimenez E et al. Isolation of commensal bacteria from umbilical cord blood of healthy neonates born by cesarean section. Curr Microbiol. 2005;51(4):270–274. 10.1007/s00284-005-0020-3

33. Louis P, Flint HJ. Formation of propionate and butyrate by the human colonic microbiota. Environ Microbiol. 2017;19(1):29–41. 10.1111/1462-2920.13589

34. Koh A, De Vadder F, Kovatcheva-Datchary P, Backhed F. From dietary fiber to host physiology: Short-chain fatty acids as key bacterial metabolites. Cell. 2016;165(6):1332–1345. 10.1016/j.cell.2016.05.041

35. Plaza-Diaz J, Ruiz-Ojeda FJ, Gil-Campos M, Gil A. Immune-mediated mechanisms of action of probiotics and synbiotics in treating pediatric intestinal diseases. Nutrients. 2018;10(1):10.3390/nu10010042

36. Mennigen R et al. Probiotic mixture vsl#3 protects the epithelial barrier by maintaining tight junction protein expression and preventing apoptosis in a murine model of colitis. Am J Physiol Gastrointest Liver Physiol. 2009;296(5):G1140–1149. 10.1152/ajpgi.90534.2008

37. Bao J et al. Dose-dependent inhibitory effect of probiotic lactobacillus plantarum on streptococcus mutans-candida albicans cross-kingdom microorganisms. Pathogens. 2023;12(6):10.3390/pathogens12060848

38. Fang SB et al. Dose-dependent effect of lactobacillus rhamnosus on quantitative reduction of faecal rotavirus shedding in children. J Trop Pediatr. 2009;55(5):297–301. 10.1093/tropej/fmp001

39. Dimidi E, Christodoulides S, Scott SM, Whelan K. Mechanisms of action of probiotics and the gastrointestinal microbiota on gut motility and constipation. Adv Nutr. 2017;8(3):484–494. 10.3945/an.116.014407

40. Mu Q, Tavella VJ, Luo XM. Role of lactobacillus reuteri in human health and diseases. Front Microbiol. 2018;9(757. 10.3389/fmicb.2018.00757

41. Thomas CM, Versalovic J. Probiotics-host communication: Modulation of signaling pathways in the intestine. Gut Microbes. 2010;1(3):148–163. 10.4161/gmic.1.3.11712

42. Young NM et al. Structural characterization and mhcii-dependent immunological properties of the zwitterionic o-chain antigen of morganella morganii. Glycobiology. 2011;21(10):1266–1276. 10.1093/glycob/cwr018

43. Monteleone G et al. Bacteria and mucosal immunity. Dig Liver Dis. 2006;38 Suppl 2(S256–260. 10.1016/S1590-8658(07)60005-X

44. Braegger C et al. Supplementation of infant formula with probiotics and/or prebiotics: A systematic review and comment by the espghan committee on nutrition. J Pediatr Gastroenterol Nutr. 2011;52(2):238–250. 10.1097/MPG.0b013e3181fb9e80

45. Qian J, Fang Z, Chang S, Zeng Z, Zhang J. Effectiveness and safety study of formula containing probiotics, prebiotics, synbiotics on fullterm infants’ growth - a systematic review and meta-analysis of randomized controlled study. Eur J Clin Nutr. 2025;79(6):504–511. 10.1038/s41430-024-01506-9

46. Underwood MA. Probiotics and the prevention of necrotizing enterocolitis. J Pediatr Surg. 2019;54(3):405–412. 10.1016/j.jpedsurg.2018.08.055

47. Perez C. [probiotics for the treating acute diarrhea and preventing antibiotic-associated diarrhea in children]. Nutr Hosp. 2015;31 Suppl 1(64–67. 10.3305/nh.2015.31.sup1.8709

48. Brandtzaeg P. Secretory iga: Designed for anti-microbial defense. Front Immunol. 2013;4(222. 10.3389/fimmu.2013.00222

49. Fadlallah J et al. Synergistic convergence of microbiota-specific systemic igg and secretory iga. J Allergy Clin Immunol. 2019;143(4):1575–1585 e1574. 10.1016/j.jaci.2018.09.036

50. Gopalakrishna KP et al. Maternal iga protects against the development of necrotizing enterocolitis in preterm infants. Nat Med. 2019;25(7):1110–1115. 10.1038/s41591-019-0480-9

